# Metabolic multi-stability and hysteresis in a model aerobe-anaerobe microbiome community

**DOI:** 10.1101/2020.02.28.968941

**Authors:** Tahmineh Khazaei, Rory L. Williams, Said R. Bogatyrev, John C. Doyle, Christopher S. Henry, Rustem F. Ismagilov

## Abstract

Changes in the composition of the human microbiome are associated with health and disease. Some microbiome states persist in seemingly unfavorable conditions, e.g., the proliferation of aerobe-anaerobe communities in oxygen-exposed environments in wounds or small intestinal bacterial overgrowth. However, it remains unclear how different stable microbiome states can exist under the same conditions, or why some states persist under seemingly unfavorable conditions. Here, using two microbes relevant to the human microbiome, we combine genome-scale mathematical modeling, bioreactor experiments, transcriptomics, and dynamical systems theory, to show that multi-stability and hysteresis (MSH) is a mechanism that can describe the shift from an aerobe-dominated state to a resilient, paradoxically persistent aerobe-anaerobe state. We examine the impact of changing oxygen and nutrient regimes and identify factors, including changes in metabolism and gene expression, that lead to MSH. When analyzing the transitions between the two states in this system, the familiar conceptual connection between causation and correlation is broken and MSH must be used to interpret the dynamics. Using MSH to analyze microbiome dynamics will improve our conceptual understanding of the stability of microbiome states and the transitions among microbiome states.

**One sentence summary:** Multi-stability and hysteresis (MSH) is a potential mechanism to describe shifts to and persistence of aerobe-anaerobe communities in the microbiome.

## Introduction

Recent evidence shows that changes in the species composition and abundance of the human microbiome can be associated with health and disease (*1-3*). Understanding the mechanisms that cause compositional shifts in healthy microbiomes, which otherwise can be remarkably stable, is challenging due to the inherent complexity of these ecosystems. A perplexing feature of some of these disturbed ecosystems is the persistence of a new microbiome state, even in seemingly unfavorable conditions. For example, in small intestinal bacterial overgrowth (SIBO), strict anaerobes that are typically found only in the colon become prominent in the small intestine and, paradoxically, persist in this environment exposed to oxygen flux from the tissue (*4, 5*). Similarly, in periodontal diseases (*6*) and in wound infections, anaerobes proliferate in oxygen-exposed environments.

One potential mechanism to explain microbiome shifts and their persistence is multi-stability (*7-13*), the concept that several steady states can exist for an identical set of system parameters (Fig. 1).

**Fig. 1.**
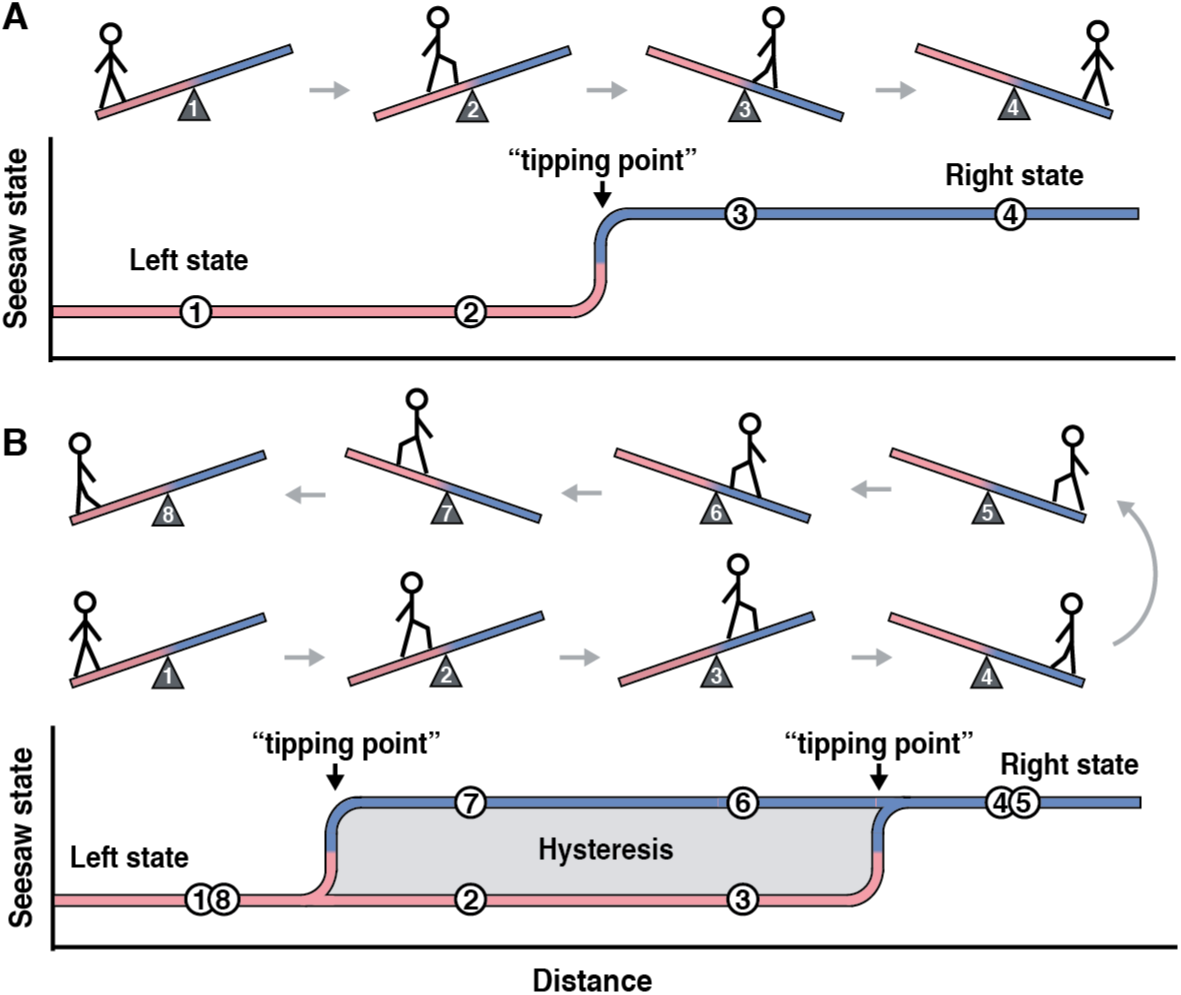
Cartoon explanation of multi-stability without hysteresis (A) and with hysteresis (B) using a seesaw and child analogy. **(A)** Consider an idealized seesaw. When a child stands on the left side of the seesaw (positions 1, 2), the seesaw is tilted to the left (the “left” state). When the child walks past the midpoint (positions 3, 4), this idealized seesaw would tilt to the right (the “right” state). The seesaw is a multi-stable system that can be in either of the two states depending on the position of the child. Moreover, the state of this seesaw is correlated with the position of the child. **(B)** Hysteresis arises if the seesaw is rusted and “sticky.” As the child walks through positions 1 and 2 past the midpoint to position 3, the seesaw remains stuck in the left-state; the child has to walk much further, to position 4, to switch it to the right state. As the child walks back to the left, through positions 5, 6, and past the midpoint to position 7, the seesaw remains stuck in the right state and it only switches back to the left state when the child reaches position 8. In the region of positions 2, 3, 6, and 7, the system exhibits multi-stability and hysteresis. Under MSH there is no correlation between the causal factor (position of the child) and the observed state of the seesaw. Furthermore, the causal factor (position of the child) cannot be used to control the state of the seesaw within the region of MSH. The system must escape the region of MSH to switch the state of the seesaw. Applying the cartoon to this manuscript, the states of the model microbiome become analogous to the states of the seesaw, and the changes in the various inputs, such as carbon and oxygen, become analogous to the changes in the child’s position.

Multi-stable systems have been described in the context of ecosystems (*14-17*) and gene-regulatory networks (*18-20*). Now, with the expanding characterization of the microbiome, there are signs that multi-stability may also exist in these communities (*21-27*). For example, compositional changes in gut microbiota are implicated in inflammatory bowel disease (*28*) and obesity (*29*). Bimodal species abundance (i.e., when a microbial species is present at either high or low levels) has been interpreted as multi-stability (*30*); however, as discussed by Gonze et al., bimodality is insufficient to prove multi-stability (*7, 31*). Some multi-stable systems can additionally exhibit hysteresis, where in response to a perturbation, a system gets “stuck” in a new steady state and the former state cannot be regained by simply reversing the perturbation (*7*). The presence of hysteresis could be hypothesized from studies of the microbiome (*32*). For example, antibiotic exposures can change the microbiome composition, and have lasting effects even after removal of the antibiotic (*33, 34*). However, it has not been rigorously tested whether multi-stability and hysteresis (MSH) can arise in a microbiome-relevant community and by what mechanism.

Here, we investigate MSH in a minimally “complex” two-species system to represent the paradoxical aerobe–anaerobe microbiome communities that persist in oxygen-exposed environments. We used two organisms prevalent in SIBO (*35*): the anaerobe *Bacteroides thetaiotaomicron* (*Bt*) that breaks down complex carbohydrates (e.g. dextran) into simple sugars and short chain fatty acids (*36*), and the facultative anaerobe (hereafter referred to as an aerobe) *Klebsiella pneumoniae* (*Kp*) capable of consuming oxygen, simple sugars and short chain fatty acids, and performing anaerobic respiration in the absence of oxygen (*37*).

## Results

To simulate how the interplay between environmental perturbations and inter-species metabolic interactions could lead to multi-stability, we first built a mathematical model. We used the dynamic multi-species metabolic modeling (DMMM) framework (*38*) to model a community of *Kp* and *Bt* in a continuously stirred tank reactor (CSTR) (Fig. 2A) with continuous input flows of dextran minimal media and varying input glucose or varying input oxygen levels (depending on the simulation). The outflow rate is equal to the inflow rate to maintain a constant reactor volume. The DMMM framework uses dynamic flux balance analysis (dFBA) (*34*), which allows us to capture temporal changes in intracellular flux rates (using the genome-scale metabolic model for each species), extracellular metabolite concentrations, and species concentrations.

**Fig. 2.**
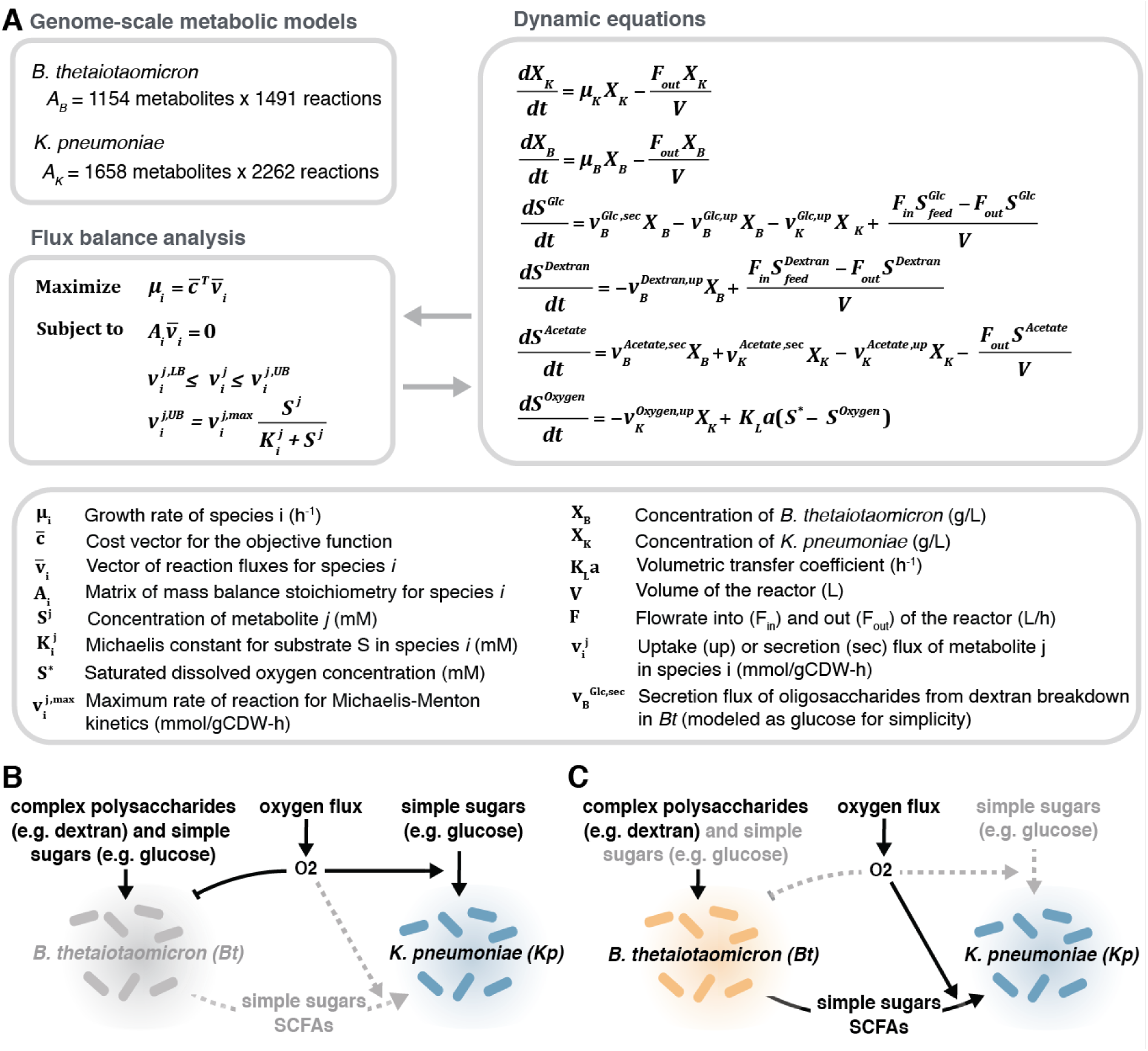
A multi-stable model system consisting of *Klebsiella pneumoniae (Kp)*, a facultative anaerobe, and *Bacteroides thetaiotaomicron (Bt)*, an anaerobe, that is relevant to the human gut microbiome. (**A**) Dynamic equations describing the model system can be solved with dynamic flux-balance analysis utilizing each species’ genome-scale metabolic model. (**B**) In the *Kp-*only state, *Bt* does not grow and *Kp* utilizes external sugars and short chain fatty acids. (**C**) In the *Kp–Bt* state, *Bt* can grow and break down complex polysaccharides into simple sugars and short chain fatty acids, which *Kp* can utilize to maintain reduced oxygen levels favorable for *Bt* growth.

Next, to computationally test whether a nutrient perturbation could lead to a change in community state, we altered glucose input concentrations (Fig. 3A), while keeping constant all other system parameters, including continuous oxygen input and continuous dextran input. The model predicted that for glucose concentrations of 0.25–3 mM in the input feed (at a constant flow rate of 0.7 mL/min for all conditions), the output state consisted solely of *Kp*, which we refer to as the *Kp*-only state (Fig. 2B). Stoichiometrically, at these glucose concentrations oxygen was not completely consumed, thus the environment was unfavorable for *Bt* growth. However, when we increased glucose input concentration to 3.25 mM, we observed a shift to a new steady state (Fig. 3A). At this “tipping point,” the environment became sufficiently anaerobic to support the growth of *Bt*. We refer to this second distinct steady state as the *Kp–Bt* (aerobe–anaerobe) state (Fig. 2C). In the *Kp– Bt* state, the *Kp* population is no longer carbon limited due to the additional carbon sources generated from the metabolism of dextran by *Bt*. Thus, *Kp* can now consume all of the available oxygen to oxidize both glucose and the additional carbon sources, resulting in anaerobic conditions. Surprisingly, this *Kp–Bt* state persisted even when we systematically reversed the input of glucose below 3.25 mM, even to 0 mM. Thus, this system shows hysteresis and multi-stability: under identical input conditions of glucose and oxygen, the system can be in either of the two possible states. We then identified tipping points for population shifts in response to input oxygen variations—with glucose kept constant (Fig. 3B). In addition to glucose input, all other parameters, including continuous dextran input, are held constant. We considered oxygen as a parameter because in host settings oxygen availability can be affected by perfusion rate, blood flow rate, immune consumption, etc. We found that we could return the system to the *Kp*-only state by increasing oxygen levels, a state switch that was not possible by manipulating glucose concentration alone. Finally, we simulated changes in both glucose and oxygen levels and characterized the landscape of multi-stability and mono-stability in the model microbial community (Fig. 3C). These simulation results illustrate that even a minimal model of microbiome with co-dependence (*39, 40*) can demonstrate dramatic MSH.

**Fig. 3.**
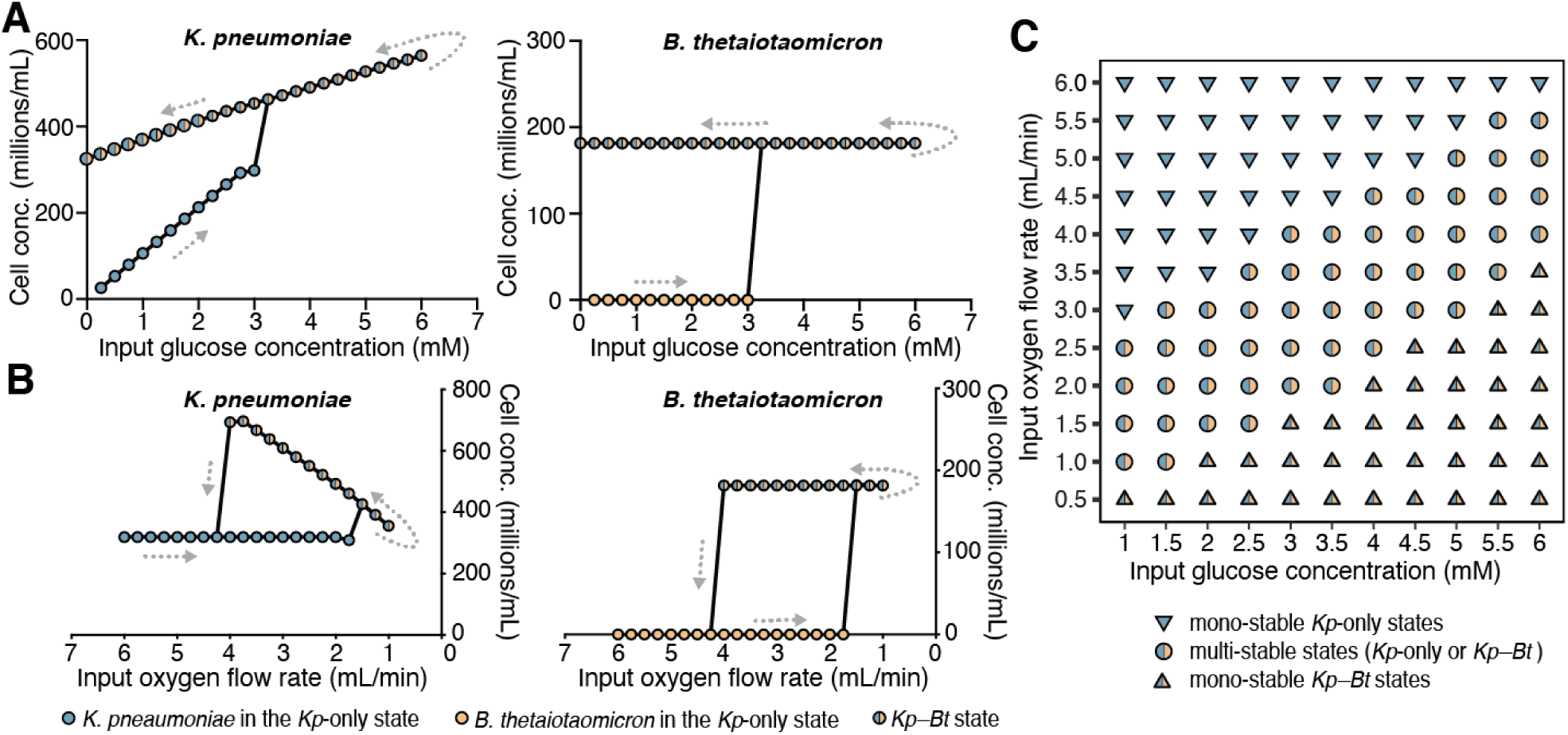
Simulations illustrating multi-stability and hysteresis in the microbial community with respect to environmental perturbations. Cell concentrations as a factor of (**A**) glucose-concentration variations in the input feed under constant input oxygen flow rate (1.7 mL/min), and (**B**) input oxygen flow variations under constant glucose concentrations (3 mM) in the input feed. Each point represents the steady-state concentration for the given species in the community after a 50-h simulation. (**C**) Regions of stability as a function of glucose concentrations in the input feed and oxygen flow rates into the reactor. In regions of multi-stability (circles) the community can exist in either a *Kp*-only state or a *Kp–Bt* (aerobe–anaerobe) state under the same conditions. In regions of mono-stability (triangles) it is only possible for one state to exist for the given set of parameters; down-pointed triangles represent mono-stable regions where *Kp*-only state exists and up-pointed triangles represent mono-stable regions where only the *Kp–Bt* state can exist.

We next tested these computational predictions experimentally in a CSTR (200 mL culture volume), and further explored the metabolic factors behind the dynamics of this aerobe–anaerobe community. We varied glucose concentrations (while keeping all other parameters constant, including continuous dextran and oxygen) and measured the steady state output composition of the microbial community by qPCR (and digital PCR; Fig. S6). Oxygen was sparged into the reactor at 3.4% of the gas feed (50 mL/min total gas feed) and kept constant for all conditions. For each steady-state condition, we collected three CSTR samples separated by at least one residence time (5 h) (Fig. S1 contains the experimental workflow).

As predicted by the mathematical models, we observed both multi-stability and hysteresis (Fig. 4A) experimentally. At 0.25 mM, 1 mM, and 2 mM glucose concentrations, the steady-state community consisted only of *Kp*; *Bt* was washed out under these conditions (Fig. 4B). The dissolved-oxygen measurements (Fig. 4C) confirmed that oxygen was not limiting under the selected parameter conditions, resulting in an aerobic environment unsuitable for *Bt* growth. As in the simulations, at 5 mM glucose, a new distinct steady state was reached where *Bt* grew in the presence of *Kp.* Although there was continuous oxygen flux into the reactor, the concentration of dissolved oxygen measured in the reactor was near zero. Next, to test for hysteresis, we reduced the glucose input back down to 2 mM, 1 mM, 0.25 mM, and 0 mM and found that the aerobe–anaerobe state persisted. The persistence of the *Kp–Bt* state (instead of a return to the *Kp*-only state), qualitatively confirmed model predictions of hysteresis, and verified that this microbial community is a multi-stable system.

**Fig. 4.**
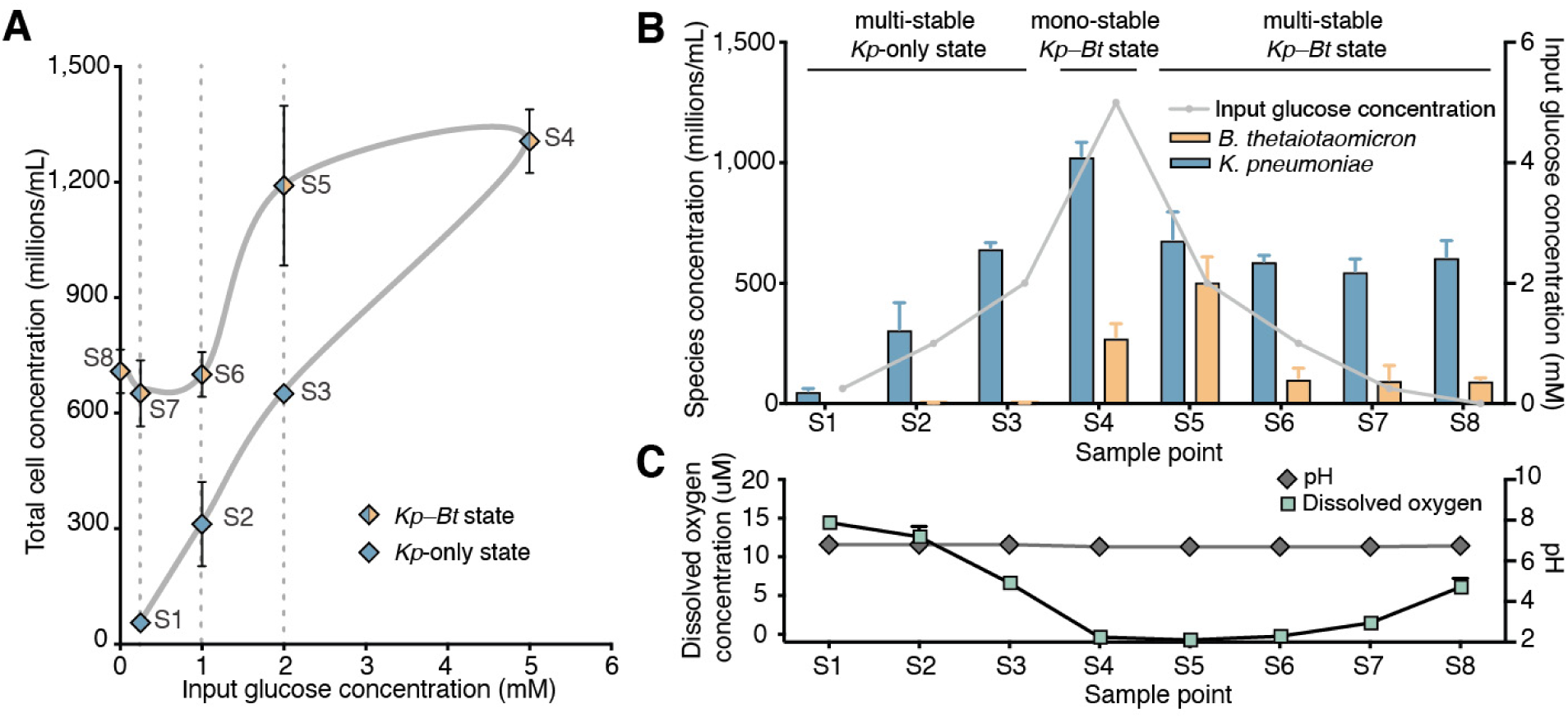
Multi-stability and hysteresis of *K. pneumoniae* (*Kp*) and *B. thetaiotaomicron* (*Bt*) community in a CSTR. (**A**) Total cell concentrations collected at the eight different steady state sample points (S1–S8) from the CSTR measured by qPCR. (**B**) Cell concentrations for each individual species in the community measured by qPCR. (**C**) pH and dissolved-oxygen concentrations measured in the CSTR for each sample point. Error bars are S.D. of at least three replicates collected at steady state (separated by >1 residence time of 5 h) from the CSTR for each of the eight steady-state glucose conditions. See Fig. S1 for bioreactor workflow.

The CSTR results demonstrate metabolic coupling and co-dependence between these two bacterial species with respect to carbon and oxygen. At sample point 8, there is no glucose input to the reactor, yet *Kp* continued to grow, indicating that *Kp* was completely dependent on *Bt* for its carbon supply. At sample point 4, *Bt* started to grow, despite the continuous oxygen input, indicating that *Bt* was dependent on removal of oxygen by *Kp*. At sample points 7 (0.25 mM glucose) and 8 (0 mM glucose), *Bt* continued to grow, despite dissolved-oxygen measurements indicating oxygen concentrations above the tolerance for *Bt* growth (Fig. 4C). This observation differed slightly from the model, suggesting that there may be additional biological factors beyond metabolic coupling and stoichiometric balance of carbon and oxygen that can affect multi-stability. Imaging revealed that in the *Kp–Bt* state bacterial aggregates were larger at lower glucose concentrations. Furthermore, fluorescent *in situ* hybridization (FISH) showed these aggregates contained both *Kp* and *Bt* (Fig. S2). We hypothesize that co-aggregation is one potential mechanism that could extend the region of hysteresis by providing microenvironments more favorable for *Bt* growth by further facilitating metabolic coupling between the two species, as observed in biofilms (*6*). Other factors, such as adhesion to the walls of the vessel may also contribute to extending the region of hysteresis.

Gene-expression analysis of CSTR samples revealed that multi-stability also occurs at the transcriptome level in both the community and in individual species. Principal component analysis (PCA) of the community-level gene expression data showed that samples clustered based on the steady state (*Kp-*only vs. *Kp–Bt)* from which they were collected (Fig. 5A). Strong clustering at the community level is expected because *Bt* is absent from the *Kp*-only state. However, when we evaluated the gene-expression profile of *Kp* (Fig. 5B), which is present in all steady state conditions, we also found clustering based on the state of the community.

**Fig. 5.**
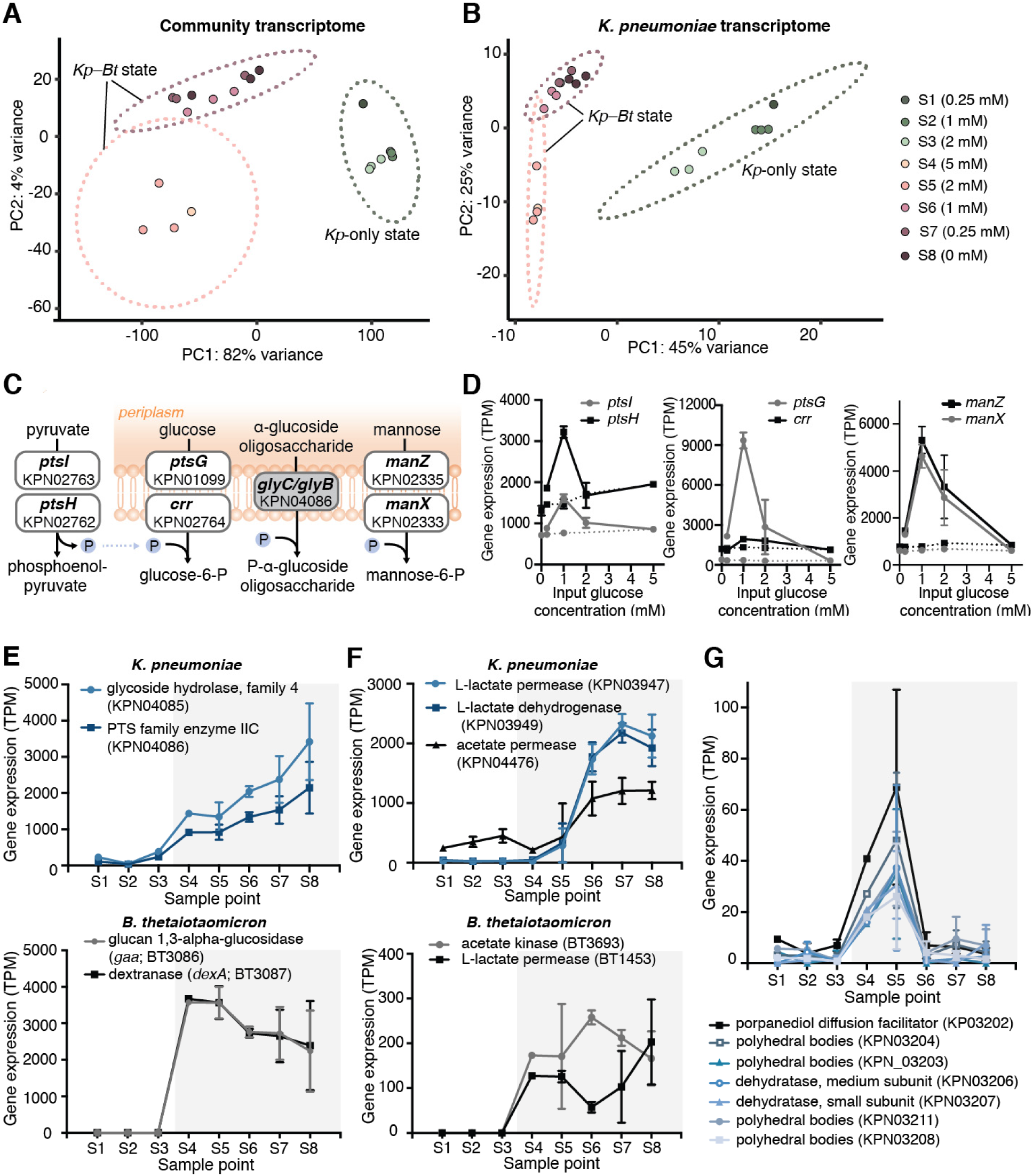
Gene-expression analysis of CSTR steady-state samples. (**A**) PCA of the community transcriptome; each point represents the combined transcriptome of *K. pneumoniae* (*Kp*) and *B. thetaiotaomicron* (*Bt*) for each sample (S1-S8). (**B**) PCA of the *Kp* transcriptome. (**C**) The most differentially regulated pathway between the *Kp-*only and the *Kp– Bt* states is the phosphotransferase system (PTS); the grey box indicates the upregulated gene; white boxes are downregulated genes. (**D**) PTS genes downregulated in the *Kp–Bt* state in *Kp*. Solid lines represent the *Kp-*only state and dashed lines represent the *Kp–Bt* state. (**E**) Gene expression, in transcripts per million (TPM), of oligosaccharide uptake in *Kp* and dextran metabolism to oligosaccharides in *Bt* for each steady-state sample point. (**F**) Expression of genes involved in acetate and lactate utilization in *Kp*, and acetate and lactate production in *Bt* for each CSTR sample. (**G**) Expression of the propanediol-utilization pathway in *Kp.* (**E–G**) Unshaded regions are the *Kp-*only state; the gray shaded region is the *Kp–Bt* state. Error bars are S.D. of three replicates collected (separated by >1 residence time) from the CSTR for each of the eight steady-state glucose conditions.

To further evaluate the proposed metabolic mechanism responsible for MSH (Fig. 2B,C), we compared metabolic regulation in *Kp* in the *Kp–Bt* state and the *Kp-*only state. We used a method from the Neilsen lab (*41*) to collect topological information from the genome-scale metabolic models and combine it with gene-expression data to identify reporter metabolites that maximally differ between the two states. Among the top reporter metabolites were pyruvate, phosphoenolpyruvate, glucose, and glucose-6-phosphate (Table S3), suggesting that the phosphotransferase system (PTS), which is involved in sugar transport, is upregulated in the *Kp*– only state relative to the *Kp-Bt* state (Fig. 5C–D). In the *Kp–Bt* state, genes involved in the alpha-glucoside linked substrates were upregulated (Fig. 5E), suggesting that *Kp* obtains some of its carbon source from oligosaccharides. These oligosaccharides are released into the environment by *Bt* through the breakdown of dextran by dextranase, an extracellular endohydrolase (*42*). *Bt* utilizes these oligosaccharides by hydrolyzing them using glucan-1,3-alpha-glucosidases. As expected, both dextranase (*dexA*) and glucan-1,3-alpha-glucosidase (*gaa*) were found to be highly expressed in *Bt* in the *Kp–Bt* state (Fig. 5E).

Our analysis (Fig. 5F) also suggested an upregulation of acetate utilization by *Kp* in the *Kp– Bt* state as inferred from the upregulation of acetate permease. Additionally, *Kp* genes involved in lactate utilization were upregulated in the *Kp–Bt* state. Upon oxygen exposure, *Bt* is known to produce lactate (*43*). A pilot experiment showed that the addition of a mixture of lactate and acetate can cause a direct “jump” from the *Kp-*only state to the *Kp–Bt* state (Fig. S5), emphasizing that short-chain fatty acids are involved in the metabolic coupling between *Kp* and *Bt* in MSH. Multi-stability of gene expression extended to the anaerobic metabolic pathway for propanediol utilization in *Kp* (Fig. 5G) (*44*). We thus infer that a subpopulation of *Kp* was undergoing anaerobic metabolism in samples 4 and 5 (of the *Kp–Bt* state), where the dissolved oxygen concentrations in the reactor were lowest (Fig. 4C). Overall, these results were consistent with the basic mechanism for MSH (Figure 2B–C) and reveal that MSH extends to the expression of genes and pathways involved in metabolic coupling between the species.

## Discussion

In this work, we used genome-scale mathematical modeling, bioreactor experiments, transcriptomics, and dynamical systems theory to show that MSH is a mechanism that can describe shifts and persistence of a two-member model microbiome aerobe–anaerobe community under seemingly paradoxical conditions (e.g., oxygen-exposed environments). We further identified key metabolic pathways involved in MSH in the *Kp-Bt* system. Future gene knock-out studies would further confirm the critical metabolic pathways responsible for MSH. A limitation of this study is the long timeframe of the CSTR experiment (Fig. S1) and that the data reported come from a single run; however, a shorter pilot experiment (Fig. S9) demonstrated similar dynamics. More broadly, identifying and interpreting MSH in human microbiomes and microbiome-associated diseases would require carefully designed longitudinal measurements and models that take into account the full complexity of microbiomes, their spatial structure, and host responses. If MSH is found, it would have profound conceptual impact. To understand and control microbial communities without MSH, one currently relies on a well-established conceptual connection between correlation, causation, and control. Consider points S1-S3 (Fig. 4). The levels of *Kp* correlate with the input glucose concentration—from a known input glucose concentration, one can infer a steady-state *Kp* concentration and vice versa. Input glucose concentration is the causal factor and therefore it can be used to control the steady-state levels of *Kp*. If MSH is identified in microbiomes, it would break this familiar conceptual connection between causation and correlation. Consider the region of hysteresis (points S1-S3 and S5-S7, Fig. 4). The observed steady-state levels of *Kp* no longer correlate with the input glucose concentration. At 2 mM input glucose, the system could be in either the *Kp*-only state S3 or the *Kp–Bt* state S5. At ∼650×10^6^ CFU/mL of *Kp*, the input glucose levels could be either 0.25 mM or 2 mM. Although there is no correlation between species’ abundance and input glucose concentration (in other words, knowing glucose concentration is not sufficient for predicting species abundance; instead, a system’s history must also be known), input glucose concentration remains the causal factor. Furthermore, under MSH, establishing causation is insufficient for achieving control: although input glucose concentration is the causal factor responsible for changes in the community state, it cannot be used to fully control the community (i.e. one cannot use changes in glucose inputs to revert the *Kp–Bt* state back to the *Kp*-only state). Alternative control strategies (e.g. changes in oxygen levels or disruption of metabolic coupling), derived from appropriate models, would need to be deployed under MSH. Therefore, recognizing whether and when MSH exists in human microbiomes will be critical for interpreting correlation and causation, and for designing therapeutic control strategies that can steer microbial communities to desirable states.

## Materials and Methods

### Model development

For the computational simulations, we used the dynamic multispecies metabolic modeling (DMMM) framework (*38*), which is an extension of dynamic flux balance analysis (dFBA) applied to microbial communities. In brief, the DMMM framework has two components: **(Component 1)** external differential equations that describe mass balances for species and metabolite concentrations in the CSTR (shown in Fig 2A and described in the SI). Unlike the traditional method for solving differential equations in a bacterial system, we do not assume that parameters such as growth rates and metabolic flux rates are constant. Instead, we allow the parameters to be dynamic because we are studying a system with potentially rich dynamics. In order to find the values for these dynamic parameters we use flux balance analysis (**Component 2**) to solve for the parameter values at every time step of the simulated time period. Component 2 includes the genome-scale models for each species. These models are used to perform flux balance analysis at every time point to obtain updated parameters for the differential equations in Component 1.

**Component 1:** The system is described as a continuous stirred tank reactor (CSTR) with the following mathematical formulation:

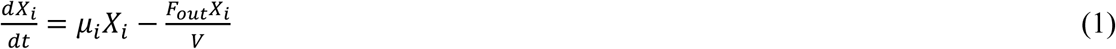

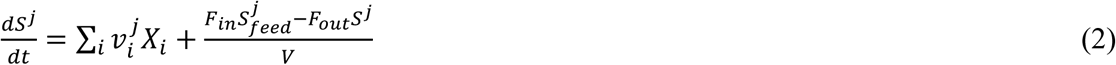

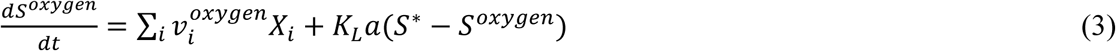

Here, V, is the volume of the reactor (constant), *X*_*i*_ is the biomass (g/L) of the i^th^ microbial species. *S*^*j*^ is the concentration (mM) of the j^th^ metabolite, *F*_*in*_ is the rate of flow (L/h) into the reactor, *F*_*out*_ is the rate of flow (L/h) out of the reactor, 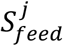 is the concentration of the j^th^ metabolite in the feed stream, *μ*_*i*_ (h^-1^) is the growth rate of the i^th^ microbial species, 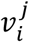 is the metabolic flux of the j^th^ substrate in the i^th^ microbial species, *K*_*L*_*a* is the volumetric oxygen transfer coefficient, and S* is the dissolved oxygen saturation concentration.

In continuous culture, substrate utilization can deviate from diauxic growth (as typically observed in batch culture) and co-substrate utilization is possible (*45*). The set of differential equations are solved using the following analytical approximation:

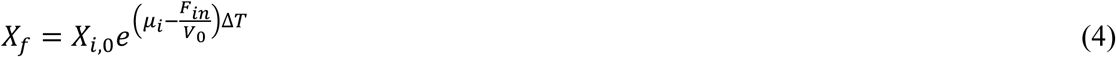

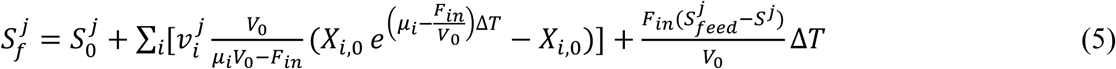

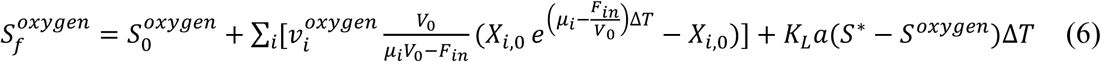

At the beginning of every time step (Δ*T*), the parameters *μ*_*i*_ and 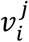 are calculated using flux balance analysis (FBA) from genome-scale models (Component 2) and fed back into equations (4), (5), and (6). This process is repeated for all time intervals in the simulated time period.

**Component 2:** Genome-scale metabolic models are used to establish genotype-phenotype relationships and capture the metabolic capabilities of each model organism. Furthermore, these models allow us to integrate metabolic network topology information with RNA sequencing data for transcriptomic analysis of the CSTR experiments (described further in “RNA Sequencing and Analysis” section). We used the published iYL1228 model of *Klebsiella pneumoniae (Kp)* MGH 78578 (*37*) and the published iAH991 model of *Bacteroides thetaiotaomicron (Bt)* VPI 5482 (*36*). Because these models are well validated (and curated), we added the minimum number of parameters that allow for integration of these models to the community dynamic flux balance analysis framework. Our changes include adding a pathway for dextran uptake and hydrolysis to glucose in the *Bt* iAH991 model. The pathway lumps hydrolysis of dextran to glucose into a single reaction. In this lumped reaction we assume that 50% of the glucose produced from dextran by *Bt* can be released into the environment for shared use. For the purpose of the simulations, dextran is assumed to be 100 glucose units. The genome-scale models are solved separately for each species by flux balance analysis (FBA) (*46*) at each time point:

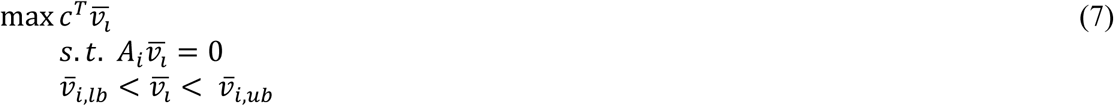

Where c is the cost vector, 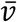 is the vector of fluxes, and *A* is the matrix of mass balance stoichiometries. The optimization criterion is biomass growth rate (for each species). For the bounds for the fluxes, we used the values in the curated, published genome-scale models (*Kp* iYL1228 and *Bt* iAH99). The uptake fluxes explicitly modeled in Component 1 (dextran, glucose, acetate, oxygen) are bounded by Michaelis–Menten kinetics:

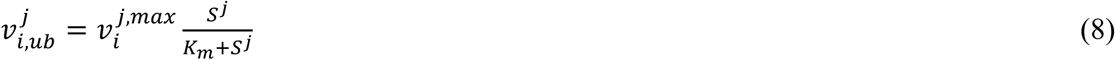

The values for 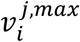 and *K*_*m*_ for some of the metabolites in the model were estimated from batch experiments. Batch culture experiments were carried out in a 96-well flat-bottom plate. Overnight cultures grown anaerobically in minimal medium supplemented with either 0.5% w/v dextran or 0.5% w/v glucose were diluted 1:20 (for *Bt*) and 1:100 (for *Kp*), and outgrown to mid-log phase. The cultures were then pelleted and re-suspended at OD 1 (for *Bt*) and OD 0.1 (for *Kp*) in carbon-free minimal medium. We added 10 µL of cells to 200 µL of minimal medium containing various concentrations (0.125 – 0.5% w/v) of the carbon source. The plate was incubated at 37 °C and OD600 was measured every 10 min. For batch cultures, Monod growth kinetics was assumed:

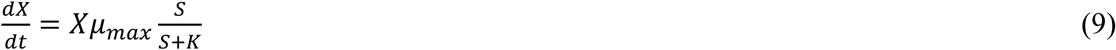

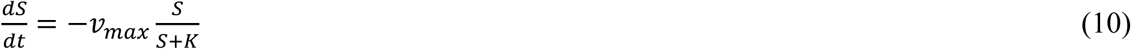

Where *μ*_*max*_ is the maximum growth rate the given bacteria can achieve on a carbon source when it is not resource limited. Growth data from replicate wells of multiple concentrations of carbon source were fitted simultaneously using Bayesian parameter estimation implemented with Markov chain Monte Carlo (MCMC) (*47*). Individual growth curves were allowed to have distinct initial cell concentrations and background values, with other parameters held constant. The fitted parameters are presented in Table S1 and Fig S3. For all other metabolites captured in the differential equations, the K_m_ and v_max_ values are assumed to be the same (v_max_ of 10 mmol/gCDW×h and K_m_ of 0.01 mM), based on literature for *Escherichia coli* (*48, 49*).

Values for parameters and initial conditions used in the model are presented in Table S2. Initial conditions are chosen to represent the experimental setup, whereby we first establish a steady state for *Kp* in the CSTR before inoculating *Bt*. Therefore, in the models we start with a higher concentration of *Kp* than *Bt*. The initial conditions for *Kp* in the reactor is arbitrarily chosen to be the experimentally measured mono-culture steady concentration of *Kp* at an input glucose concentration of 0.25 mM. The initial conditions for *Bt* in the reactor is 0.0015 g/L, which is equivalent to addition of 1 mL of OD 1 *Bt* into the reactor, as done experimentally. For most steady state conditions, glucose is limiting and therefore the initial conditions for glucose concentration in the reactor is chosen to be 0mM.

We note that although an individual FBA simulation (Component 1) is linear, a dynamic FBA (the combination of Component 1 and Component 2) is not linear. The behavior of the cells will shift dramatically (a nonlinear response) when substrate levels in the environment cross certain thresholds. The two biggest sources of nonlinearity in our model system are binary growth/no-growth behavior of *Bt* in the presence or absence of O_2_, and the binary capacity of *Kp* to utilize carbon from dextran (through its breakdown into oligosaccharides by *Bt*) in the presence or absence of *Bt*. To model the binary growth/no-growth behavior of *Bt* in the presence of O_2_, we included a conditional operator before the FBA simulations for each time step: If the O_2_ levels are above 350 nM, the growth rate of *Bt* is set to zero and the FBA is not run for *Bt*. Whereas if O_2_ levels are lower than 350 nM O_2_, then the FBA simulation for *Bt* is allowed to proceed. The value for the O_2_ growth threshold for the model is arbitrarily chosen from the concentration range reported for the aerotolerant *Bt* (*50*).

To computationally identify the regions of stability with respect to glucose and oxygen (Fig. 3C and Fig. S4), we varied oxygen input flow rates at constant input glucose concentration for each glucose condition examined. We evaluated 11 glucose conditions ranging from 1 mM to 6 mM. For each given glucose input concentration, we started with oxygen at an input flow rate of 6 mL/min and ran the simulation for 50 h to ensure the system reached a steady state. We then decreased the oxygen input by 0.5 mL/min intervals down to 0.5 mL/min; for each oxygen condition, ensuring the system reached a steady state (we refer to the 6mL/min - 0.5mL/min oxygen variations for a given constant glucose input concentration as the “forward simulations”). The concentration of oxygen input at which *Bt* starts to grow is identified as the “tipping point” to the mono-stable *Kp– Bt* state. After running the 0.5mL/min oxygen simulation we increased the concentration back to 6 mL/min at intervals of 0.5 mL/min (we refer to the 0.5 mL/min - 6 mL/min oxygen variations as the “reverse simulations”). The concentration of oxygen at which *Bt* can no longer grow and gets washed out is identified as “tipping point” to the mono-stable *Kp-*only state. The region between these two tipping points to the mono-stable *Kp–Bt* state in the forward simulations and the monostable *Kp*-only state in the reverse simulations is identified as the region of bi-stability. The colors in Fig. S4 represent the steady state concentration of *Kp* in the “reverse simulations” divided by the steady state concentration of *Kp* in the “forward simulations.” In regions of mono-stability the concentration of *Kp* is the similar in both the “forward” and “reverse simulations”, and therefore has a value of approximately 1.

### Continuous culture of *K. pneumoniae* and *B. thetaiotaomicron*

Continuous culture experiments were carried out in a 500 mL bioreactor (Mini-bio Applikon Biotechnology, Delft, Netherlands) with a total culture volume of 200 mL. Minimal media (3.85 g/L KH_2_PO_4_, 12.48 g/L K_2_HPO_4_, 1.125 g/L (NH_4_)_2_SO_4_, 1X MMS (20X MMS: 17.6 g/L NaCl, 0.4 g/L CaCl_2_, 0.4 g/L MgCl_2_×6H2O, 0.2 g/L MnCl_2_×4H2O, 0.2 g/L CoCl_2_×6H2O), 10 mL/L Wolfe’s mineral solution (*51*), 10 mL/L Wolfe’s vitamin solution (*51*), 4.17 µM FeSO4×7H20, 0.25 mM cysteine, 1 µM menadione, 2 µM resazurin, 1 g/L dextran (Sigma D5376, avg. mol. wt 1.5e6-2e6) and glucose at varying concentrations) was purged with 100% N_2_, stored under anaerobic conditions prior to use, and maintained under N_2_ during operation of the CSTR. We calibrated the dissolved oxygen probe by aerating the reactor with CO_2_ (5 ml/min) and N_2_ (45 ml/min). The stable measurement without O_2_ input was taken to be 0 µM dissolved oxygen. We then sparged the reactor with O_2_ (1.7 ml/min), CO_2_ (5ml/min), and N_2_ (43.3 ml/min). This stable measurement was taken to be 30.714 µM O_2_ (calculated assuming 6.056 mg/L dissolved oxygen at 1 atm and 37 °C).

For the experiments, the bioreactor was sparged with 50 mL/min total gas (1.7 mL/min O_2_, 5 mL/min CO_2_, and balance of N_2_), and agitated with two six-bladed Rushton turbines operated at 750 rpm. Temperature was maintained at 37°C, and a residence time of 5 h (input and output flow rates of 40 ml/h flowrate) was used for all experiments. Dissolved oxygen, pH, and biomass were monitored throughout. For initial inoculation of *Kp*, 1 mL of OD 1 culture was injected through the septum, and grown in batch culture until stationary phase (indicated by a levelled biomass readout and an increase of dissolved oxygen levels) before beginning continuous culture. For every experimental condition examined (input glucose concentration) in the CSTR, we waited for the system to first reach steady state (at least 24 h). To assess whether a steady state had been reached, we monitored the total biomass in the reactor using a real-time OD probe. Once the system reached steady state, we took three samples over the course of 24-48 h. The time interval between each sample collection was at least one residence time (5 h). Residence time is defined as the time it takes to entirely exchange the volume of the reactor. For introduction of *Bt*, a log phase (OD 0.6-0.8) anaerobic culture grown in minimal media with 0.5% dextran and 2 mM cysteine was pelleted (5 min at 3500 g) and washed twice using dextran/glucose-free anaerobic minimal media. Cells were carbon-starved at 37 °C for 30 min, washed (once), and re-suspended in dextran/glucose-free minimal media to OD 1. We used 1 mL of this *Bt* cell suspension for inoculation into the reactor and a sample was collected immediately after inoculation. A subsequent sample was collected for quantification after at least 2 residence times had passed. In the *Kp*-only state conditions (0.25 mM, 1 mM, 2 mM glucose), *Bt* is washed out, as described in the results section. To ensure reproducibility of a washout for these conditions, the *Bt* inoculation and sample collection process was repeated a total of three times. In *Kp–Bt* state conditions (5 mM, 2 mM, 1 mM, 0.25 mM, and 0 mM glucose), where *Bt* growth persisted, re-inoculation of *Bt* was no longer necessary for each new glucose steady state condition; three samples separated by at least one residence time were collected for each steady state condition. To collect samples, ∼0.5 mL of culture was removed from the bioreactor in a 3 mL luer-lock syringe and discarded before collection of 1.5–2 mL culture. Supernatant from 700 µL of the collected sample was stored at −80 °C for SCFA analysis, a 50 µL sub-sample was treated with DNAse (2.5 µL of NEB DNase I 2000 u/mL per 50 µL) for subsequent DNA extraction, and two 250 µL aliquots were used for extraction of RNA.

### Quantification of bacterial abundance

CSTR culture samples were treated with NEB DNAse I (100 u/mL final concentration) for 10 min at 37 °C immediately after collection. DNA was extracted using the ZyGEM prepGEM Bacteria kit (ZyGEM, Southampton, England) according to the manufacturer protocol. Samples were extracted in 100 uL total volume (20 µL culture sample and 80 µL of extraction mixture), incubated at 37 °C for 15 min, 75 °C for 5 min, 95 °C for 5 min, then cooled to 4 °C. DNA was stabilized by adding 10X TE to a final concentration of 1X TE before storage at 4 °C.

### qPCR quantification

Extracted DNA was quantified by qPCR using the Eco Real-time PCR system (Illumina, San Diego, CA, USA). The components in the qPCR mix used in this study were as follows: 1 µL of extracted DNA, 1X SsoFast™ EvaGreen Supermix (Bio-Rad Laboratories, Hercules, CA, USA), 500 nM forward primer, and 500 nM reverse primer. For detection of each bacterial species in the community primer sets specific to *Bt* (forward primer: 5′-GGAGTTTTACTTTGAATGGAC-3′; reverse primer: 5′-CTGCCCTTTTACAATGGG-3’) and *Kp* (forward primer: 5′-ATTTGAAGAGGTTGCAAACGAT-3′; reverse primer: 5′-TTCACTCTGAAGTTTTCTTGTGTT-3′) were used. Quantification of cell concentrations were determined using DNA standards of single species prepared using 10X serial dilutions of log phase cultures extracted as above. Cell concentrations of standards were determined by hemocytometer. For conversion of OD and cell concentration to biomass concentration (gram cell dry weight/L), 100 mL of culture for each individual species incubated anaerobically at 37 °C was harvested and pellets were dried at 80°C for ∼48 h before recording mass.

### Digital PCR quantification

Archived DNA samples from the CSTR were quantified by digital PCR (dPCR) using a QX200 Droplet Digital PCR System (Bio-Rad). The components in the dPCR mix were as follows: 1 µL of dilutions of extracted DNA, 1X QX200 ddPCR EvaGreen Supermix (Bio-Rad), 500 nM forward primer, and 500 nM reverse primer. For detection of each bacterial species in the community primer sets specific to *Bt* (forward primer:5′-GGTGTCGGCTTAAGTGCCAT-3′; reverse primer: 5′-CGGAYGTAAGGGCCGTGC-3′) and *Kp* (forward primer:5′-ATGGCTGTCGTCAGCTCGT-3′; reverse primer: 5′-CCTACTTCTTTTGCAACCCACTC-3′) were used. The dPCR mix was loaded into DG8 Cartridges (Bio-Rad), which were filled with QX200 Droplet Generation Oil for EvaGreen (Bio-Rad) and loaded into the QX200 Droplet Generator (Bio-Rad). The generated droplets were transferred to the C1000 Touch Thermal Cycler (Bio-Rad) for the following protocol: 5 min at 95 °C, 40 cycles of 30 sec at 95 °C, 30 sec at 60 °C (*Kp*) or 65 °C (*Bt*), and 30 sec at 72 °C (ramping rate reduced to 2°C/s), and final dye stabilization steps of 5 min at 4 °C, and 5 min at 90 °C. The stabilized plates were loaded into the QX200 Droplet Reader and analyzed using the QuantaSoft Analysis Software (Bio-Rad).

### RNA sequencing and analysis

From the CSTR samples, a 250 µL aliquot was used for metatranscriptomic analysis. The freshly collected CSTR sample was immediately placed into Qiagen RNAprotect Bacteria Reagent (Qiagen, Hilden, Germany) for RNA stabilization. RNA was extracted using the Enzymatic Lysis of Bacteria protocol of the Qiagen RNeasy Mini Kit and processed according to the manufacturer’s protocol. DNA digestion was performed during extraction using the Qiagen RNase-Free DNase Set. The quality of extracted RNA was measured using an Agilent 2200 TapeStation (Agilent, Santa Clara, CA, USA). Extracted RNA samples were prepared for sequencing using the NEBNext Ultra RNA Library Prep Kit for Illumina (New England Biolabs, Ipswitch, MA, USA) and the NEBN<u>e</u>Ext Multiplex Oligos for Illumina. Libraries were sequenced at 100 single base pair reads and a sequencing depth of 10 million reads on an Illumina HiSeq 2500 System (Illumina, San Diego, CA, USA) at the Millard and Muriel Jacobs Genetics and Genomics Laboratory, California Institute of Technology. Raw reads from the sequenced libraries were subjected to quality control to filter out low-quality reads and trim the adaptor sequences using Trimmomatic (v. 0.35). Because our samples were a mixture of *Kp* and *Bt* cells, to separate the reads for each species, we did the following: Reads that aligned to rRNA and tRNA of *Bt* and *Kp* were first removed, as those sequences contain overlapping reads between the two species. Each sample was then separately aligned to *Bt* VPI-5482 (Genome accession number: GCA_000011065.1) and *Kp* MGH-78578 (Genome accession number: GCA_000016305.1) using Bowtie2 (v. 2.2.5) and quantified using the Subread package (v. 1.5.0-p1). Gene expression was defined in transcripts per million (TPM) for each species and gene expression analysis was performed using DESeq2 (v. 1.22.2; default settings, which provides two-tailored p-values).

To determine the most differentially regulated metabolic pathway between the *Kp-Bt* and *Kp*-only states we used an approach by Patil et al. (*41*), which combines gene expression data with topological information collected from genome-scale metabolic models. In brief, in this approach the metabolic network is presented as a bipartite undirected graph, where metabolites as well as enzymes are represented as nodes (this graph is obtained from genome-scale models). Differential data can be mapped on the enzyme nodes of the graph with specification of the significance of differential gene expression for each enzyme, i. We used DESeq to perform our differential gene expression analysis between sample points in the two states (*Kp*-only and *Kp-Bt*) to obtain *P*-values. The *P*-values are subsequently converted to Z-score for an enzyme node using the inverse normal cumulative distribution. Finally, the Z-score of each metabolite node is calculated based on the normalized transcriptional response of its k neighboring enzymes:

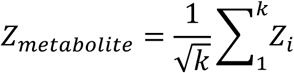

The metabolites with the highest Z-scores mark the pathways with substantial regulation between two states.

## Supplementary Materials

**Fig. S1.** The workflow for the continuous bioreactor experiment

**Fig. S2.** Imaging of samples collected from the continuously stirred tank reactor experiments.

**Fig. S3.** Bayesian parameter fitting of K and v_max_ to experimental batch growth data.

**Fig. S4.** A quantitative view of regions of stability as a function of glucose concentrations in the input feed and oxygen flow rates into the reactor.

**Fig S5.** State switch from *Kp*-only state (state A) to the *Kp–Bt* state (state B) by the transient addition (pulse) of short chain fatty acids.

**Fig S6.** Quantification of *B. thetaiotaomicron* and *K. pneumoniae* in archived CSTR samples using digital PCR

**Fig. S7.** Simulations of extracellular concentrations of dextran and glucose (A), and acetate and oxygen (B) as a factor of input glucose concentration under constant oxygen feed (flow rate of 1.7 mL/min).

**Fig. S8.** *P*-values and individual data points for (A) Fig. 3A (B) Fig. 3B (C) Fig. S5 and (D) Fig. S6 of the manuscript.

**Fig. S9.** A pilot CSTR experiment demonstrating state-switching and hysteresis.

**Table S1.** Bayesian parameter estimation for K and v_max_ used in the Michaelis–Menten equations to constrain nutrient uptake flux rates for flux balance analysis calculations.

**Table S2.** Values for the parameters used in the dynamic flux balance analysis simulations.

**Table S3.** The top scoring 50 metabolites involved in the most regulated metabolic pathways, ordered by Z-score value.

**Table S4.** *P*-values between statistically significant samples (gene expression values) for K. pneumoniae in Fig. 5.

## Acknowledgments

We thank Jens Nielsen (Chalmers University of Technology), Richard Murray (Caltech), Jared Leadbetter (Caltech), and Elaine Hsiao (UCLA) for helpful discussions. We thank Roberta Poceviciute (Caltech) for imaging samples, Kevin Winzey (Caltech) for performing digital PCR analysis of samples, and Natasha Shelby for contributions to writing and editing this manuscript.

## Funding

This work was supported in part by Army Research Office (ARO) MURI contract #W911NF-17-1-0402, NSF Emerging Frontiers in Research and Innovation (EFRI) Grant 1137089A, NSERC fellowship PGSD3-438474-2013 [to T.K.], and the Center for Environmental Microbial Interactions (CEMI). This work was also supported by the Millard & Muriel Jacobs Genetics and Genomics Laboratory at Caltech and we thank director Igor Antoshechkin for his assistance.

## Author contributions

T.K., S.R.B., J.C.D. C.S.H., and R.F.I conceptualized the study. T.K., C.S.H. and R.F.I. contributed to the computational investigation. T.K., R.L.W., and R.F.I. contributed to the experimental investigation. T.K. wrote the manuscript and all authors contributed to the final submission of the manuscript.

## Competing interests

Authors declare no competing interest.

## Data and materials availability

All associated raw sequencing data have been deposited in the Sequence Read Archive (Bio-Project Accession Number PRJNA580293). All other data will be made publicly available upon publication at CaltechDATA.

