## Supplemental Information for "Metabolic multi-stability and hysteresis in a model aerobe-anaerobe microbiome community"

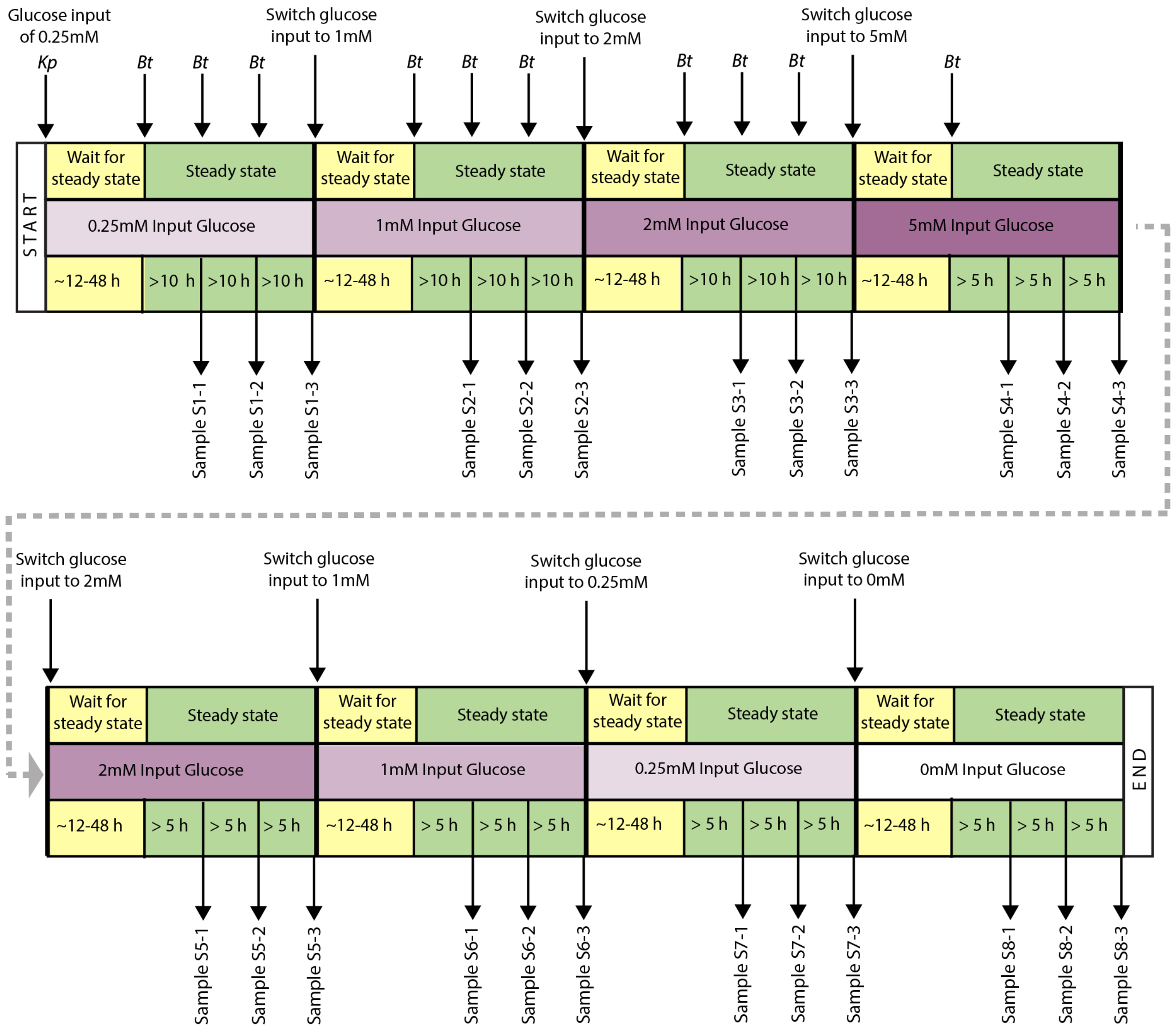


**Fig. S1. The workflow for the continuous bioreactor experiment indicating the points at which we inoculated with bacteria, took samples, and changed glucose concentrations of the input feeds.**

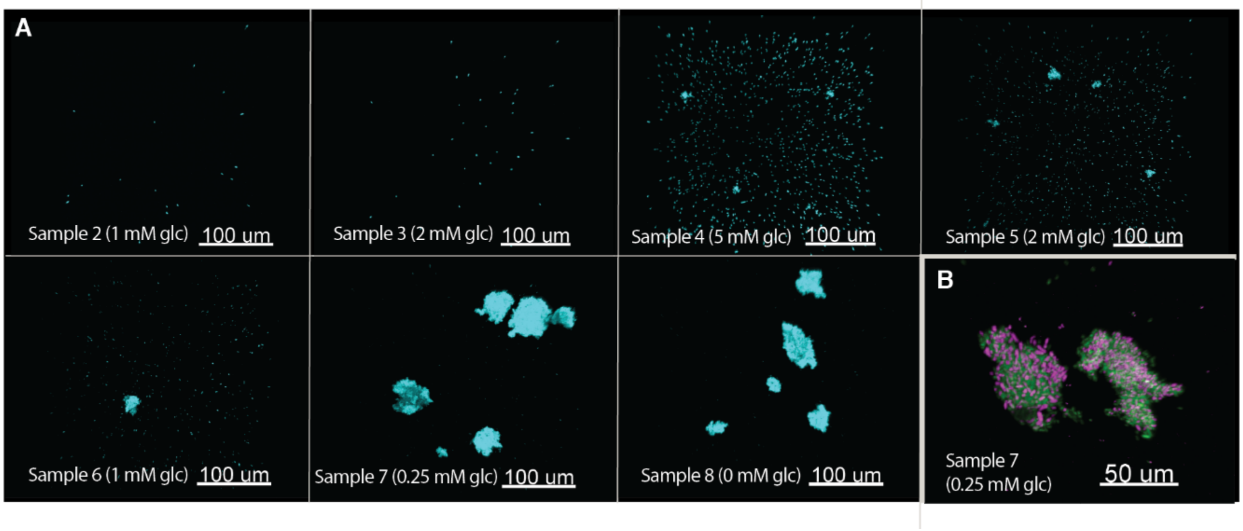


**Fig. S2. Imaging of samples collected from the continuously stirred tank reactor experiments.** (**A**) Total bacteria staining of each sample using DAPI (**B**) The species composition of aggregates using the GAM42a (green) and CFB560 probe (pink) for Gammaproteobacteria and Bacteroidetes, respectfully.

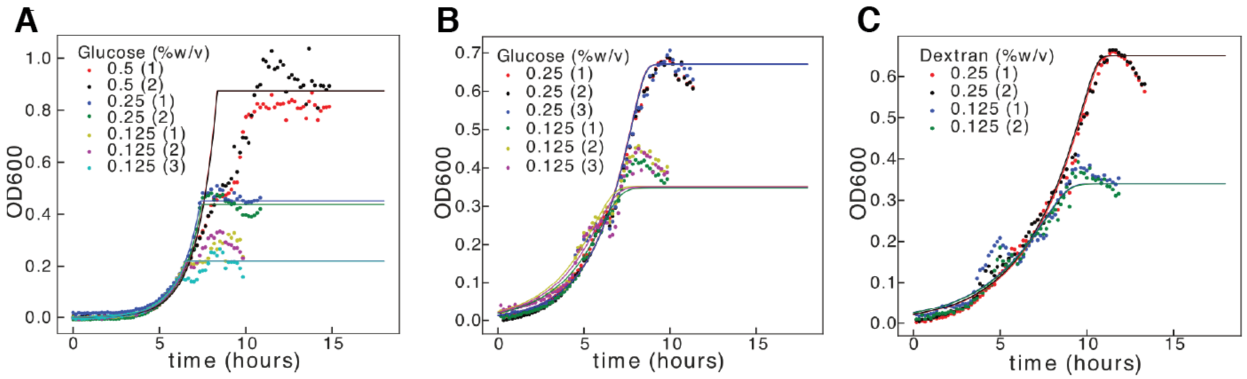


**Fig. S3.** Bayesian parameter fitting of K and $v_{max}$ to experimental batch growth data for (**A**) *Klebsiella pneumoniae* on glucose (**B**) *Bacteroides thetaiotaomicron* on glucose, and (**C**) *Bacteroides thetaiotaomicron* on dextran. Two or three technical replicates were used for each concentration of substrate examined.

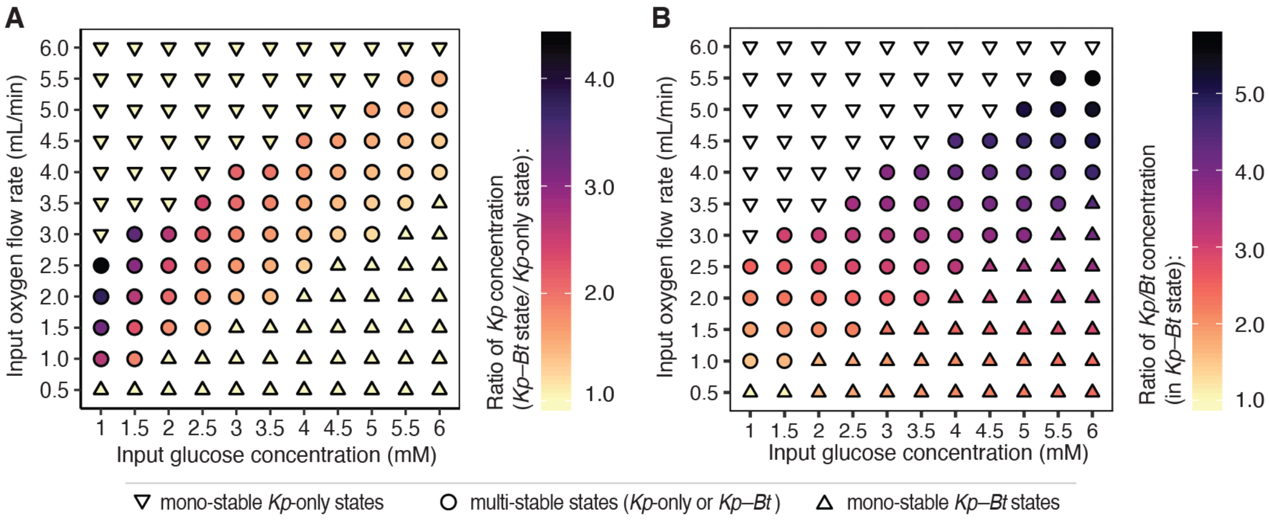


**Fig. S4**. **A quantitative view of regions of stability as a function of glucose concentrations in the input feed and oxygen flow rates into the reactor.** In regions of multi-stability (circles) the community can exist in either a *Kp*-only state or a *Kp–Bt* (aerobe–anaerobe) state under the same conditions. In regions of mono-stability (triangles) the community can only exist in either a *Kp*-only or a *Kp–Bt* state. **(A)** Deviation from yellow indicates the increase of *Kp* concentration in the *Kp–Bt* state relative to the *Kp*-only state (e.g. concentration of *Kp* in *Kp–Bt* state divided by the concentration of *Kp* in the *Kp*-only state). **(B)** Ratio of *Kp* concentration to *Bt* concentration in the *Kp–Bt* state. Ratios were not calculated for the *Kp*-only state as *Bt* does not grow in that state (white down-pointed triangles).

Proof-of-Concept Experiment for Community Control via SCFA

The goal of this proof-of-concept experiment was to show that we can jump directly from one steady state (*Kp*-only) to an alternate steady state (*Kp-Bt* state) without changing glucose concentration (i.e., without taking the system through a progression of glucose concentrations, from low to high and then back to low). We predicted that we could make this direct jump if we give the system the metabolites that are exchanged in the alternate (*Kp-Bt*) steady state. We began by flowing minimal media with a 2 mM input glucose in the CSTR. In this condition, as demonstrated in Fig. 4 (and Fig. S5), only *Kp* is able to grow at steady state and *Bt* gets washed out. We then switched the media to 2 mM glucose + 7 mM acetate + 7 mM lactate (the metabolites identified in the *Kp-Bt* state by transcriptomics analysis) and we observed that when *Bt* was inoculated, it did not wash out but grew along with *Kp*. Finally, we switched the media back to 2 mM glucose and observed that both *Bt* and *Kp* continued to grow at a steady state (Fig. S5). This experiment demonstrates another method we can use to manipulate a system that exhibits MSH. We show that we can directly provoke the system to move to a different steady state by adding the metabolites exchanged by the species.

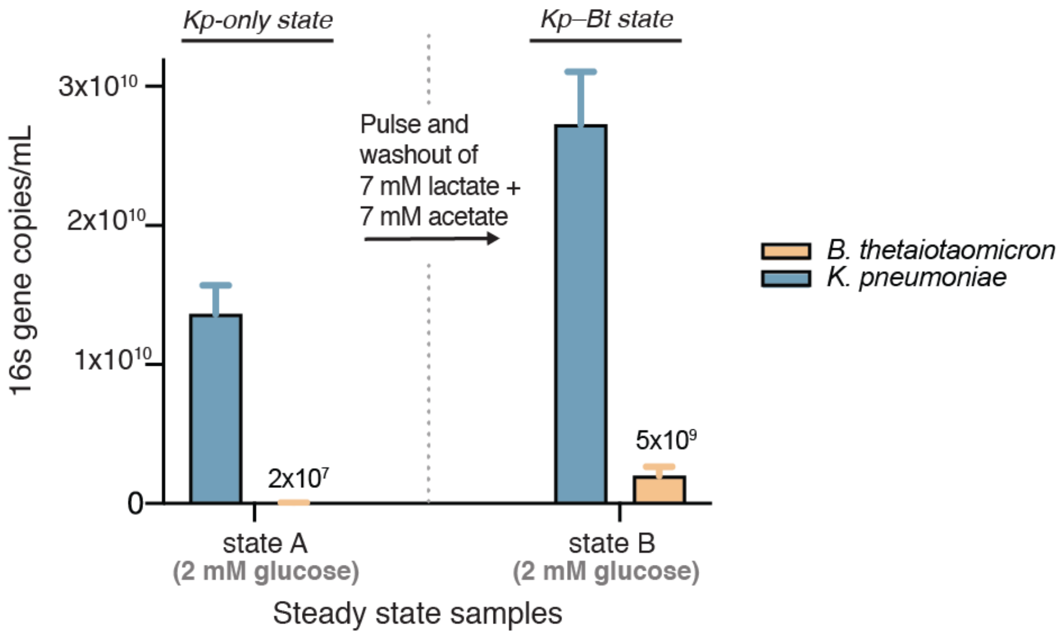


**Fig S5. State switch from *Kp*-only state (state A) to the *Kp–Bt* state (state B) by the transient addition (pulse) of short chain fatty acids (addition of 7 mM lactate + 7 mM acetate were flowed with 2 mM glucose for 24 h).** Measurements for each species are made by digital PCR using 16S primers specific to each species. Error bars are S.D. of three replicates collected from the CSTR (over 24 h after steady state was reached, each separated by more than 1 residence time) for each of the steady-state conditions (state A and state B).

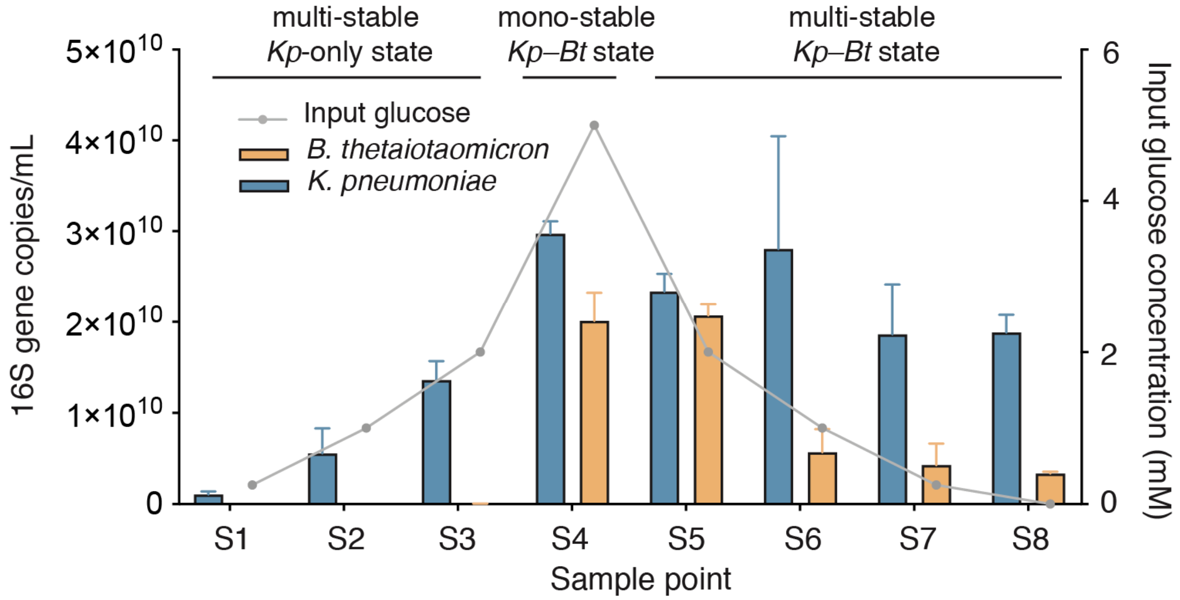


**Fig S6. Quantification of *B. thetaiotaomicron* and *K. pneumoniae* in archived CSTR samples using digital PCR.** Primers used targeted the 16S ribosomal RNA gene for each species. Error bars are S.D. of three replicates collected (separated by >1 residence time) from the CSTR for each of the eight steady-state glucose conditions.

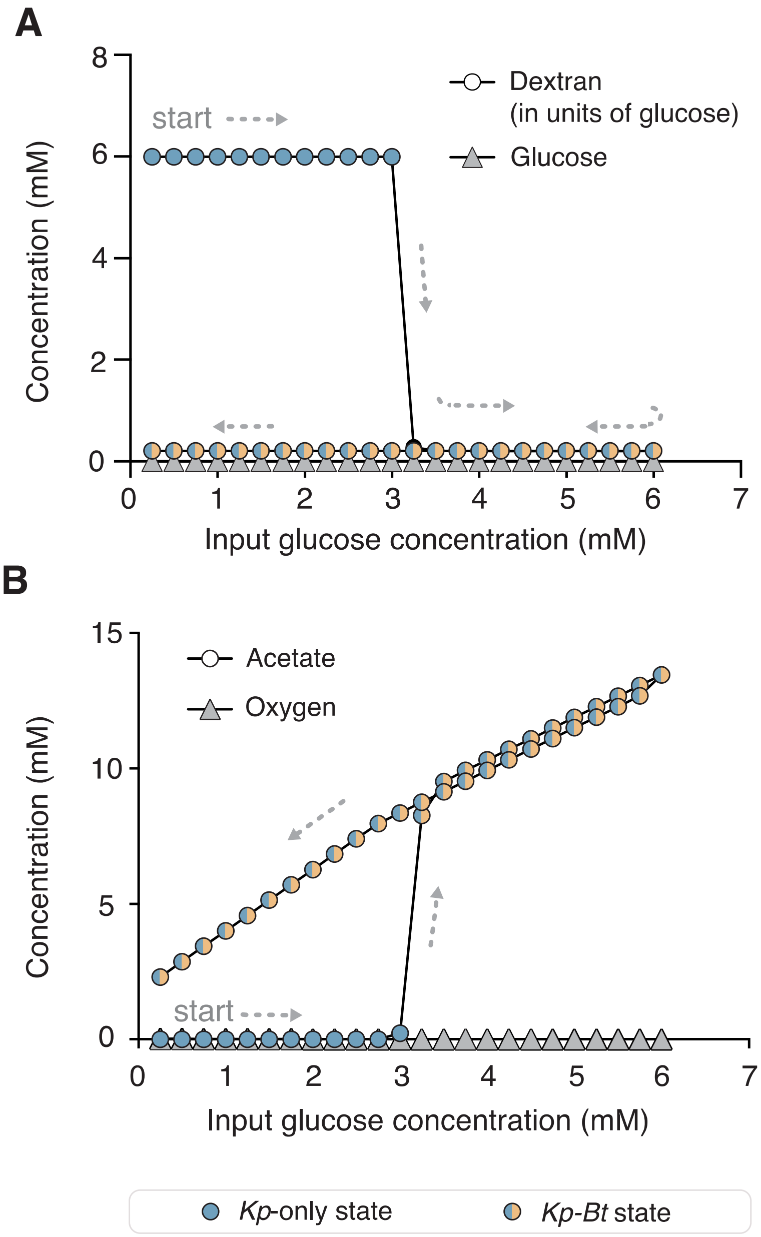


**Fig. S7. Simulations of extracellular concentrations of dextran and glucose (A), and acetate and oxygen (B) as a factor of input glucose concentration under constant oxygen feed (flow rate of 1.7 mL/min).** Each point represents the steady-state concentration after a 50-h simulation. The values for glucose (A) and oxygen (B), in both the *Kp-Bt* state and the *Kp*-only state are indicated by gray triangles. Arrows are used to indicate the direction of the changes in dextran (A) and acetate (B) concentrations as the community shifted from the *Kp*-only state (blue circles) to the *Kp-Bt* state (blue/orange circles) and then persisted in the Kp-Bt state as a function of input glucose (increased and then decreased).

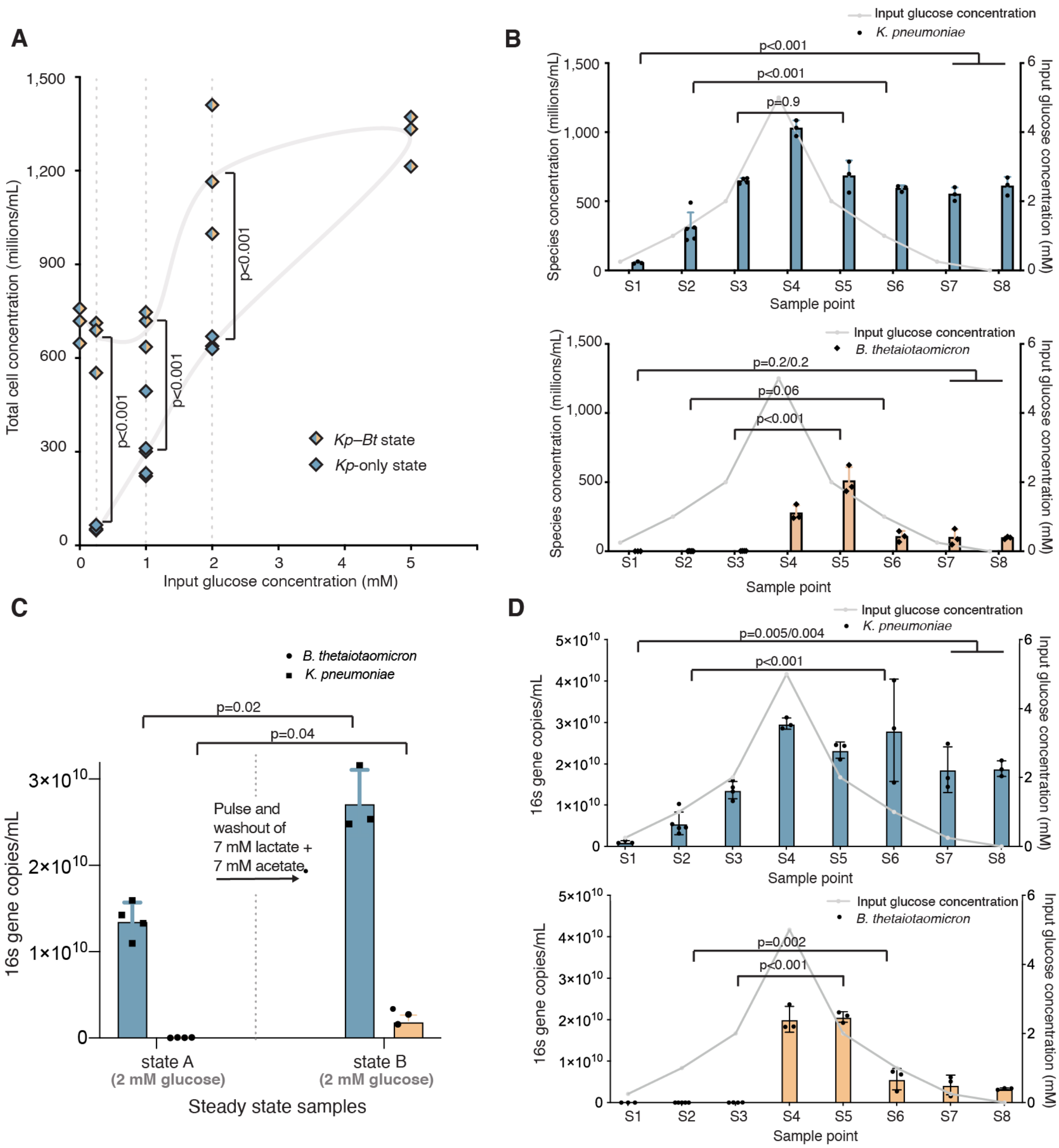


**Fig. S8. *P*-values and individual data points for (A) Fig. 4A (B) Fig. 4B (C) Fig. S5 and (D) Fig. S6 of the manuscript.** For (A) and (B) and (D) pair-wise comparisons were performed using the Tukey-Kramer test. For (C) FDR-corrected Welch's p-values were calculated.

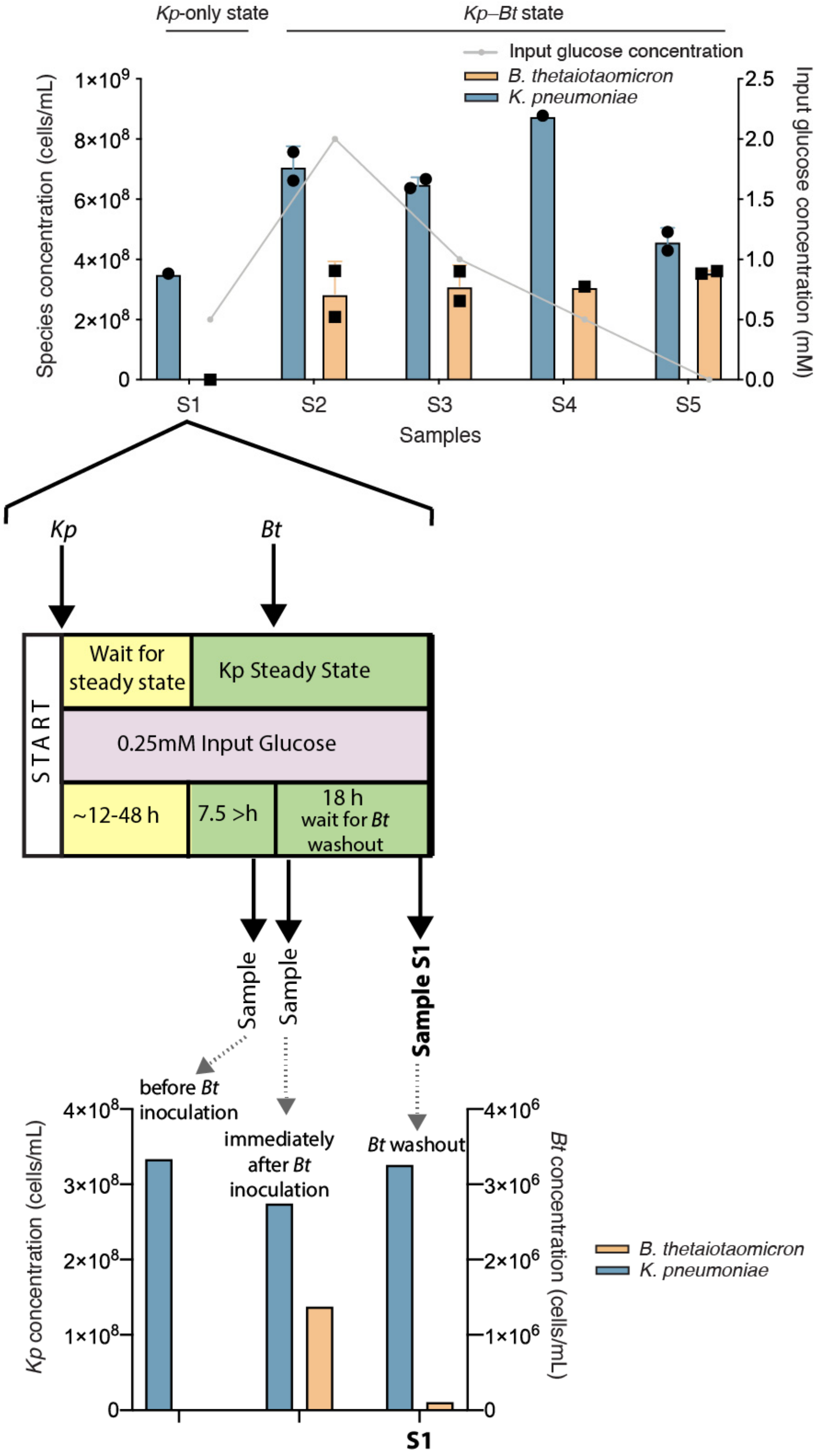


**Fig. S9. A pilot CSTR experiment demonstrating state-switching and hysteresis.** The pilot CSTR experiment differed from the main experiment in two ways: (1) mode of oxygen delivery and (2) residence time. In the manuscript (Fig. 4), oxygen was directly sparged at 1.7 mL/min, whereas in the pilot run we sparged air at 8 mL/min (equivalent to 1.68 mL/min O2). The residence time in the manuscript was 5 h whereas in the pilot it was 7.5 h. One steady-state sample was collected for samples S1 and S4 and two steady-state samples, separated by at least one residence time, were collected for samples S2, S3, and S5.

**Table S1.** Bayesian parameter estimation for K and $v_{max}$ used in the Michaelis–Menten equations to constrain nutrient uptake flux rates for flux balance analysis calculations.

| Species | Carbon Source | K (mM) | Vmax (mmol/gCDW×h) |
|---|---|---|---|
| *K. pneumoniae* | Glucose | 0.00005 | 26.1 |
| B. *thetaiotaomicron* | Glucose | 1.4 | 10.9 |
| B. *thetaiotaomicron* | Dextran | 0.008 | 0.08 |

**Table S2.** Values for the parameters used in the dynamic flux balance analysis simulations.

| **Parameter** | **Value** |
|---|---|
| Initial glucose concentration | 0 mM |
| Initial dextran concentration | 0.06 mM |
| Dissolved oxygen saturation concentration | S* = $(\frac{\text{Input O2 flowrate}}{\text{Total flowrate}})$/(O2 fraction in air) x (dissolved oxygen saturation)<br>▪ Input oxygen flow rate = varies<br>▪ Total flow rate = 50 mL/min<br>▪ Oxygen fraction in air = 0.2095<br>▪ Dissolved oxygen saturation (at 37 °C and 20 g/kg salinity) = 0.189 mM |
| Initial concentration of all other metabolites | 0 mM |
| Initial *K. pneumoniae* concentration | 0.05 g/L |
| Initial *B. thetaiotaomicron* concentration | 0.0015 g/L |
| Flow rate (into and out of the reactor) | 0.04 L/h |
| Volumetric oxygen transfer coefficient (Kla) | 47.4 $h^{-1}$ |
| Volume | 0.2 L |
| Time span (for each steady state condition) | 50 h |
| Time step (ΔT) | 0.05 h |

**Table S3.** The top scoring 50 metabolites involved in the most regulated metabolic pathways, ordered by *Z*-score value. This analysis was performed using gene expression data between samples S8 and S3.

| Metabolite name | Number of genes | Number of reactions | Z score |
|---|---|---|---|
| Pyruvate | 260 | 57 | 1063.96 |
| Phosphoenolpyruvate | 174 | 25 | 961.45 |
| H+ | 2468 | 859 | 629.11 |
| ADP | 964 | 247 | 480.86 |
| ATP | 1158 | 326 | 399.89 |
| H2O | 1786 | 523 | 350.85 |
| CO2 | 198 | 78 | 306.33 |
| Phosphate | 978 | 259 | 301.71 |
| D-Glucose | 54 | 9 | 257.35 |
| Malonyl-[acyl-carrier protein] | 50 | 16 | 219.07 |
| D-Glucose 6-phosphate | 50 | 13 | 207.81 |
| H+ | 726 | 241 | 198.26 |
| acyl carrier protein | 140 | 51 | 195.38 |
| D-Fructose 6-phosphate | 46 | 16 | 129.56 |
| D-Fructose | 26 | 3 | 109.83 |
| alpha,alpha-Trehalose 6-phosphate | 18 | 5 | 105.50 |
| N-Acetyl-D-glucosamine 6-phosphate | 20 | 4 | 95.97 |
| D-Glucosamine 6-phosphate | 18 | 7 | 88.31 |
| N-Acetyl-D-glucosamine | 22 | 2 | 87.79 |
| D-Mannose 6-phosphate | 14 | 5 | 82.90 |
| N-Acetyl-D-mannosamine 6-phosphate | 12 | 2 | 81.25 |
| D-Glucose | 60 | 20 | 77.32 |
| D-Mannose | 18 | 2 | 75.20 |
| N-Acetyl-D-mannosamine | 18 | 2 | 75.20 |
| D-Glucosamine | 18 | 2 | 75.20 |
| Maltose | 18 | 3 | 73.60 |
| Maltose | 22 | 7 | 72.65 |
| Glyceraldehyde 3-phosphate | 46 | 14 | 70.02 |
| Trehalose | 20 | 3 | 69.47 |
| Acetaldehyde | 26 | 12 | 67.28 |
| Maltohexaose | 26 | 8 | 65.21 |
| Maltopentaose | 24 | 7 | 59.01 |
| Formate | 24 | 5 | 56.09 |
| Sodium | 42 | 16 | 54.91 |
| D-Glucose 1-phosphate | 30 | 10 | 54.58 |
| Maltotetraose | 22 | 6 | 52.09 |

| | | | |
|---|---|---|---|
| N-acetylmuramate 6-phosphate | 10 | 2 | 48.06 |
| Sodium | 50 | 16 | 48.02 |
| D-Galactose | 22 | 6 | 47.82 |
| D-Galactose | 20 | 5 | 46.58 |
| L-Serine | 42 | 24 | 45.97 |
| Acetate | 42 | 17 | 44.94 |
| L-Lactate | 8 | 4 | 44.93 |
| Glycerol 3-phosphate | 42 | 15 | 44.41 |
| S-Adenosyl-L-methionine | 46 | 22 | 43.02 |
| Ubiquinone-8 | 106 | 20 | 41.72 |
| Maltose 6-phosphate | 8 | 1 | 41.35 |
| S-Adenosyl-L-homocysteine | 40 | 19 | 41.28 |
| Ubiquinol-8 | 108 | 21 | 39.82 |
| Flavin adenine dinucleotide reduced | 28 | 13 | 39.73 |

**Table S4.** *P*-values between statistically significant samples (gene expression values) for *K. pneumoniae* in Figures 5D, 5E, and 5F obtained using t-test with FDR-correction.

| **Figure** | **Gene** | ***P*-value between multi-stable samples S2 and S6** |
|---|---|---|
| **5D** | KPN_01099 (*ptsG*) | 0.0001 |
| **5D** | KPN_02333 (*manX*) | 0.0002 |
| **5D** | KPN_02335 (*manZ*) | 0.0002 |
| **5D** | KPN_02762 (*ptsH*) | 0.0002 |
| **5D** | KPN_02763 (*ptsI*) | 0.0002 |
| **5D** | KPN_02764 (*crr*) | 0.002 |
| **5D** | KPN_03949 | 0.0003 |
| **5E** | KPN_04085 | 0.0001 |
| **5E** | KPN_04086 | 0.0002 |
| **5F** | KPN_04476 | 0.01 |

### Contributions of non-corresponding authors

#### Tahmineh Khazaei

- Hypothesis ideation with SRB.
- Design of study.
- Performed preliminary experiments with SRB: evaluating *Kp-Bt* community growth under various glucose conditions in batch culture.
- Built the mathematical model used in this study (Figure 2). Using the mathematical models predicted state-switching and hysteresis within the community and identified regions of bi-stability with respect to glucose and oxygen input conditions (Figure 3).
- Designed CSTR experiments. These experiments were performed by RLW and TK.
- Established the protocol for short chain fatty acids measurements (further optimized by RLW).
- Performed qPCR of the CSTR samples (Figure S5).
- Established the protocol for RNA extraction of CSTR samples for RNA sequencing. Performed the RNA extraction of all CSTR samples for RNA sequencing.
- Established the bioinformatics pipeline for processing and analyzing the CSTR samples (mixed-species samples). Processed and analyzed the RNA sequencing data (Figure 5).
- Wrote and made figures for the manuscript.

#### Rory L. Williams

- Performed preliminary plate reader experiments testing state switching with BT/KP and BT/E. coli that determined we would use KP in CSTR experiments
- Established the CSTR workflow, optimized media conditions, and performed the CSTR experiments with help from TK (Figure 3).
- Designed CSTR experiments with TK.
- Worked with Nathan Dalleska to optimize HPLC for the measurement of SCFAs in CSTR samples.
- Performed qPCR of the CSTR samples (Figure 3).
- Characterized some of the Michaelis Menton constants used in the mathematical models. This was done through batch experiments for growth of Bt and Kp on various substrates and Bayesian parameter inference.
- Helped TK in preparing the manuscript.

#### Said R. Bogatyrev

- Hypothesis ideation with TK.
- Designed and performed preliminary experiments with TK: evaluating *Kp-Bt* community growth in batch culture as a function of substrate concentration, selectivity, and redox potential in the system.
- Performed statistical analysis
